# BCI Toolbox: An Open-Source Python Package for the Bayesian Causal Inference Model

**DOI:** 10.1101/2024.01.02.573851

**Authors:** Haocheng Zhu, Ulrik Beierholm, Ladan Shams

## Abstract

Psychological and neuroscientific research over the past two decades has shown that the Bayesian causal inference (BCI) is a potential unifying theory that can account for a wide range of perceptual and sensorimotor processes in humans. Therefore, we introduce the BCI Toolbox, a statistical and analytical tool in Python, enabling researchers to conveniently perform quantitative modeling and analysis of behavioral data. Additionally, we describe the algorithm of the BCI model and test its stability and reliability via parameter recovery. The present BCI toolbox offers a robust platform for BCI model implementation as well as a hands-on tool for learning and understanding the model, facilitating its widespread use and enabling researchers to delve into the data to uncover underlying cognitive mechanisms.

## Introduction

It has been proposed that the human brain functions like a Bayesian statistical machine (Knill & Pouget, 2004), with the nervous system continuously processing uncertain sensory information from different modalities to infer the causes of sensory observation. Bayesian causal inference (BCI) model (Körding et al., 2007) is a normative Bayesian model of this process in which an inference is made both about the causal structure (common cause vs. independent causes) as well as the sources of the sensory inputs. In the BCI model, the inference about the causal structure and the inference about the source are done in a coherent unified fashion, and involve competition between two hypotheses–were the sensory information generated by a common cause or by independent causes–to account for observed sensory measurements.

During the past two decades, the BCI model has been extended and employed in a large variety of perceptual and sensorimotor domains (for a review see, Shams & Beierholm, 2010; 2022), including temporal numerosity judgment (Rohe et al., 2019; Wozny et al., 2008), spatial localization judgment (Körding et al., 2007; Wozny & Shams, 2011; Wozny et al., 2010; Odegaard et al, 2015), size-weight illusion paradigm (Peters et al., 2016), rubber-hand illusion paradigm (Samad et al., 2015; Chancel et al., 2022; Chancel & Ehrsson, 2023), and heading perception (Dokka et al., 2019). Given this empirical evidence, the BCI model has been recognized as a potential unifying framework in neuroscience (Shams & Beierholm, 2022).

Inspired by the substantial potential of the BCI model and the increasing demand for Bayesian data analysis within the domain of neuroscience and psychology, we introduce the Bayesian causal inference toolbox (BCI Toolbox), a zero-programming software in Python to the scientific community as a tool for understanding and using the BCI model. The BCI Toolbox contains a graphical user interface (GUI) for primary use and well-studied mathematical functions for advanced use. We aim to facilitate the use of the BCI model; therefore, the present GUI includes user-friendly model fitting and simulation functionalities. The software can be installed from the online documentation (https://bcitoolboxrmd.readthedocs.io/en/latest/index.html), GitHub (https://github.com/evans1112/bcitoolbox), or via PIP (https://pypi.org/project/bcitoolbox).

Here, we provide an overview of the algorithm implementation and software architecture of the BCI Toolbox and discuss the performance of the BCI model through parameter recovery, further corroborating the reliability of the model.

### Design and implementation

In principle, the general implementation is based on the Bayesian causal inference model of multisensory perception (Körding et al., 2007). To describe the basic structure of the model, here we use the example of hearing a sound and seeing a sight and the task of estimating the location of the sound (*s_A_*). However, the model is general (not specific to any sensory modalities or perceptual tasks) and the toolbox implementation allows for combination of two sensations from any modalities, and a variety of perceptual tasks as discussed in the following sections.

Figure 1A shows the generative model of BCI, wherein two possible causal structures, namely a common cause and independent causes, can give rise to sensory inputs *x_A_* and *x_V_*. During the inference stage or perception, these two hypotheses compete to explain the sensory observations in order to estimate the perceptual variables of interest, e.g., the location of the auditory event (*s_A_)* and the location of the visual event (*s_V_*). As shown in Figure 1B, the underlying causal structure of the stimuli is inferred based on the available sensory evidence and prior knowledge. Each stimulus or event *s* in the world causes a noisy sensation *x_i_* of the event. We use the generative model to simulate experimental trials and subject responses by performing Monte Carlo simulations. Each sensation is modeled using the likelihood function *p*(*x_i_*|*s*). Trial-to-trial variability is introduced by sampling the likelihood from a normal distribution around the true locations *s_A_* and *s_V_*, plus bias terms **γ**_*A*_ and **γ**_V_ for auditory and visual modalities, respectively. This simulates the corruption of auditory and visual sensory channels by independent Gaussian noise with standard deviation *σ_A_* and *σ_V_*, respectively. In other words, the sensations *x*_A_ and *x*_V_ are simulated by sampling from the distributions shown in Eqs 1 and 2.

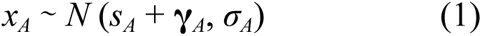

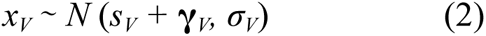

**Figure 1.**
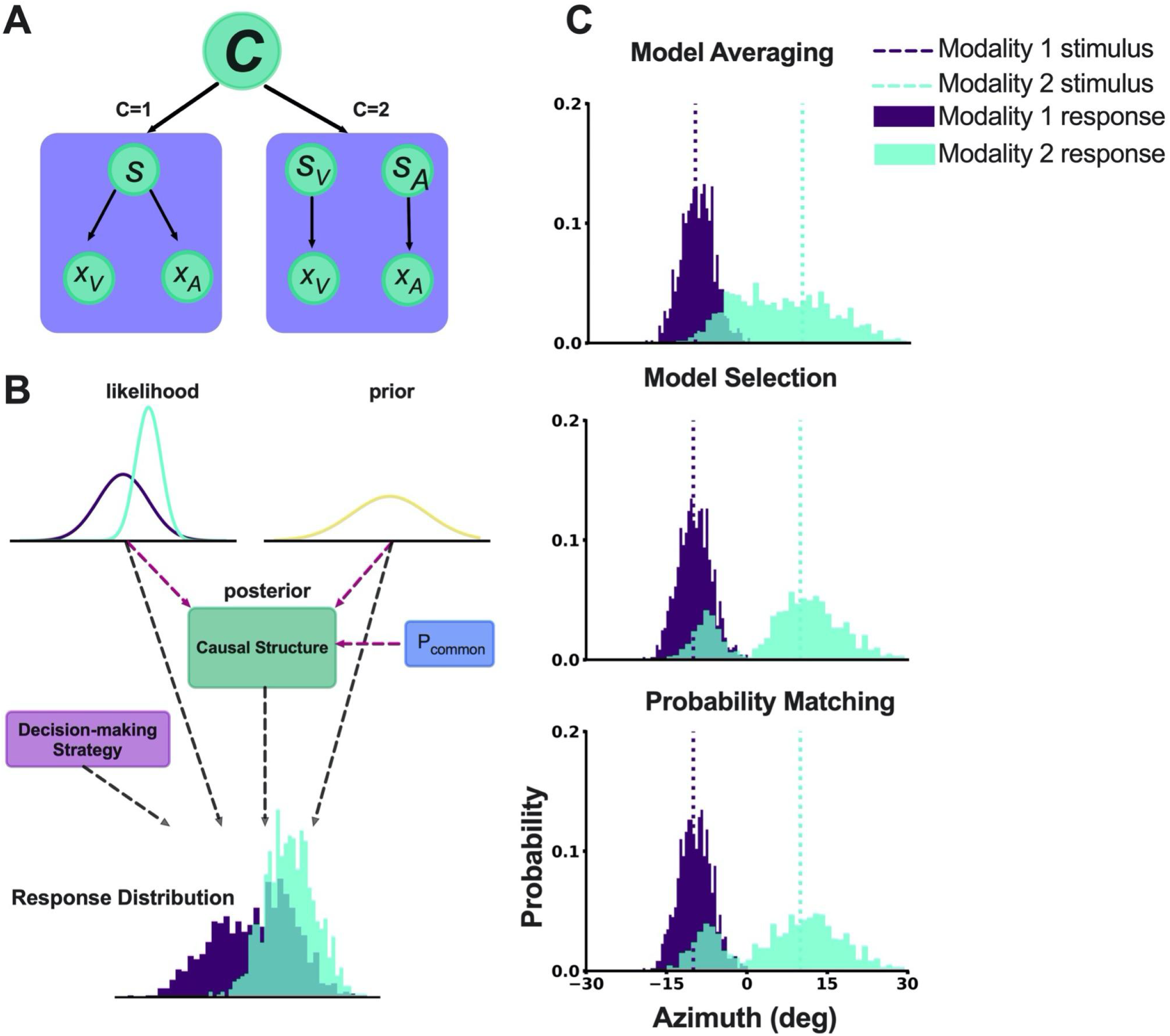
The general structure of the BCI model and simulation results of BCI Toolbox. (A) The generative model of BCI, assumes that there is either one cause (C=1) or two causes (C=2), leading to the creation of the perceptual variables (*s* or *s_A_* and *s_V_*). (B) The structure of the hierarchical BCI model in the BCI Toolbox. The causal structure is inferred by combining sensory likelihood and prior (prior stimulus expectation and P_common_). P_common_ represents *a priori* expectation of a common cause). The observer response is based on the inferred causal structure, and the decision-making strategy. (C) The one-dimensional model simulation results (generated by p_common_=0.5; **σ**_1_=3; **σ**_2_=8; **σ**_P_=30; μ_P_=0; s_1_=-10; s_2_=10) from 3 different decision-making strategy using the BCI Toolbox.

We assume there is a prior bias for the sensory information (Odegaard et al. 2019), modeled by a Gaussian distribution centered at *μ_P_*. The standard deviation of the Gaussian, *σ_P_*, determines the strength of the bias. Therefore, the prior distribution of sensory information is:

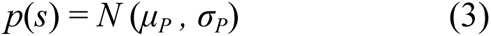

As the causal structure is unknown to the nervous system, it must be inferred using sensory information and prior knowledge. The probability of each causal structure is computed using Bayes Rule as follows:

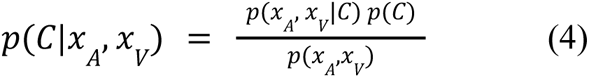

The optimal estimate of source *s* in each modality depends on the causal structure. If the sensations are produced by independent causes, the estimate of *s* is a weighted average of the unisensory signal and the prior for *s*:

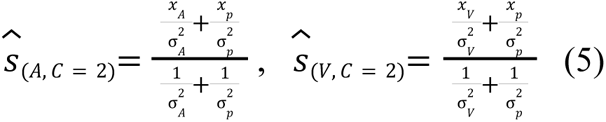

If the sensations are produced by a common cause, the estimate of *s* is a weighted average of the both sensory signals and the prior for *s*:

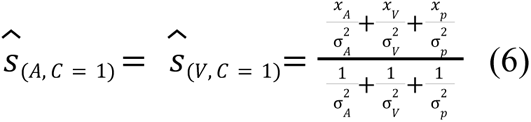

As can be seen in Eq. (4), the inference about the causal structure is probabilistic, and therefore, there is uncertainty associated with each causal structure. The optimal estimate of the sources *s_A_* and *s_V_* depend on the goal of the perceptual system in a given task. If the goal is to minimize the average error in the magnitude of the source estimates, i.e., a sum squared error cost function, then optimal strategy for achieving this goal is *model averaging*, in which optimal estimates corresponding to both causal structures are taken into account, however, proportional to their respective probability (Körding et al.2007; Wozny et al. 2010).

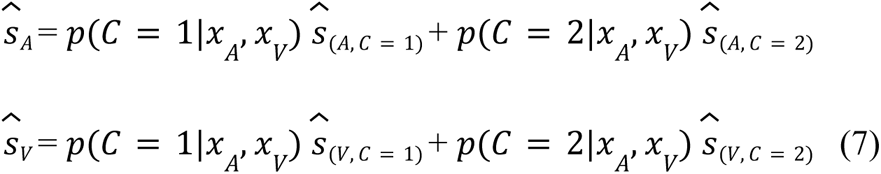

However, there are other plausible cost functions. Indeed, Wozny et al. (2010) showed that in a spatial localization task a number of observers’ performance was more consistent with *model selection* or *probability matching* strategies. If the nervous system’s goal is to minimize the error in the inference of causal structure, this would result in a decision strategy that selects the optimal estimates of sensory information based on the posterior probability of causal structure, which is referred to as *model selection* (Eq. 8).

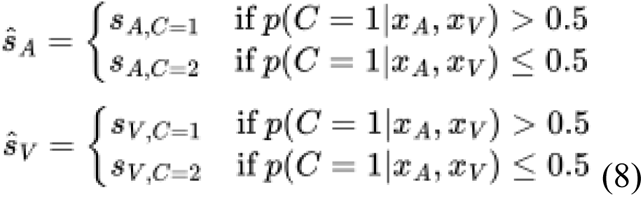

*Probability matching* is a stochastic strategy where the nervous system selects a causal structure according to its inferred probability (Eq. 9). This strategy is optimal if learning is also a factor in the utility function (Wozny et al., 2010).

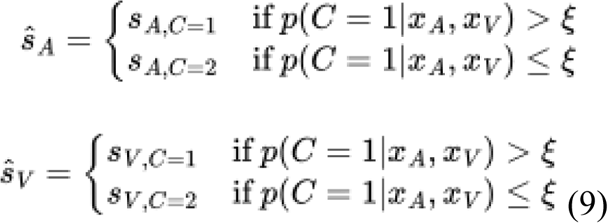

ξ ∈ [0:1] uniform distribution and sampled on each trial.

More details on the model can be found elsewhere (e.g., Körding et al., 2007; Wozny & Shams, 2011); we point the interested reader to earlier publications for additional information.

#### Graphical User Interface (GUI)

To facilitate user experience, we developed an intuitive Graphical User Interface (GUI) for researchers. Figure. 2 illustrates the overall structure of the interface. The GUI currently offers two main functions, model simulation and model fitting, which will be described in detail below.

**Figure 2.**
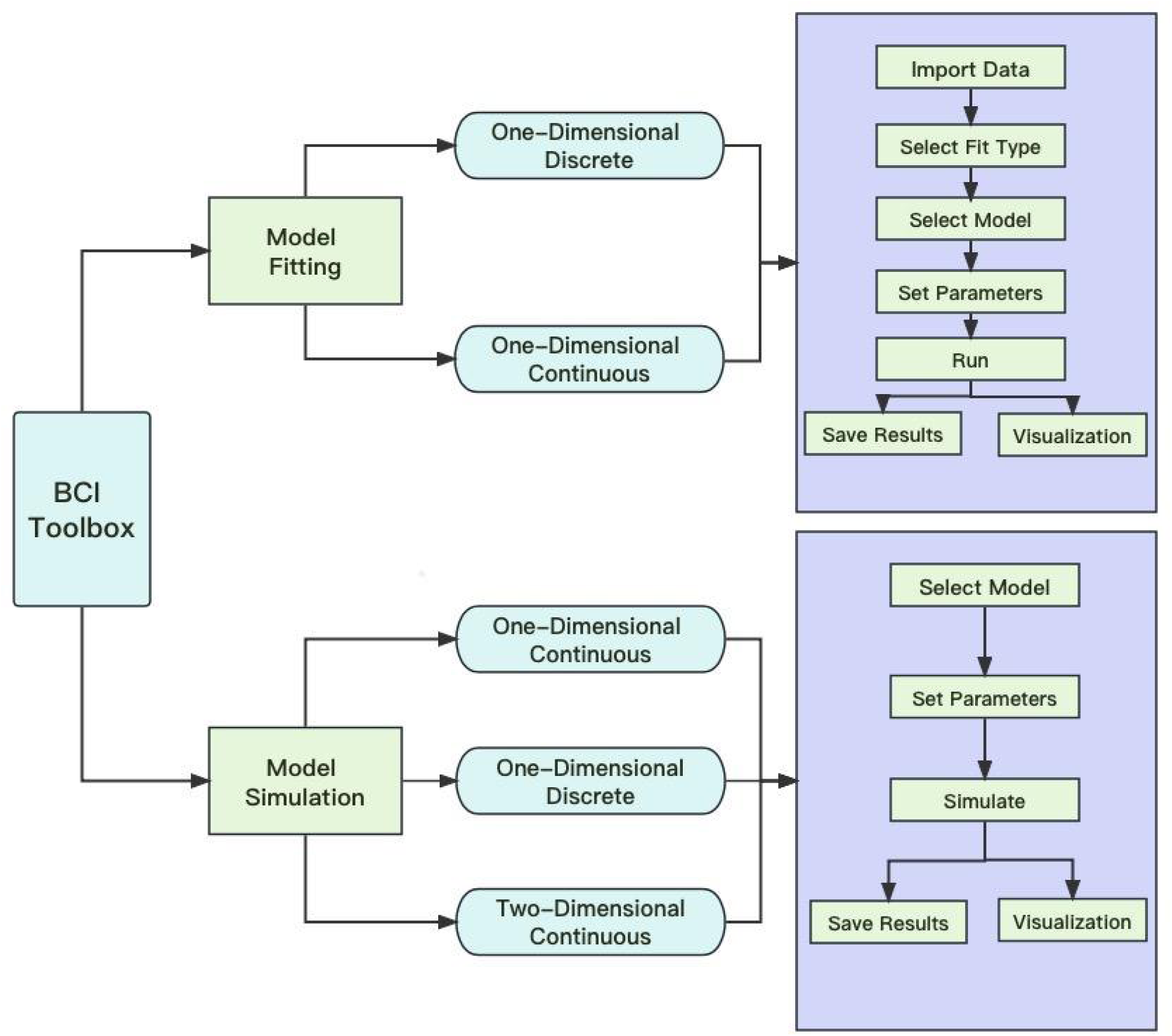
An overview of the BCI Toolbox GUI. The GUI provides two main functions: model fitting and model simulation. In the model fitting section, the GUI incorporates two data types: discrete and continuous data. In the model simulation section, the GUI incorporates one-dimensional and two-dimensional simulations. For more details, see BCI Toolbox documentation: https://bcitoolboxrmd.readthedocs.io/

#### Model Fitting

In this module, users can input behavioral data for model fitting. The BCI Toolbox provides fitting for two data types: discrete and continuous data, and can either maximise the likelihood of the data given the model, or minimise the squared error between model and data. Users have the flexibility to select various decision strategies (corresponding to different cost functions (Wozny et al. 2010)). Additionally, they can adjust seven key parameters to tailor the BCI model to their experimental paradigm. These parameters include *p_common_* (the prior expectation of a common cause), *μ_1_*, *μ_2_* (the mean of the likelihood), *σ_1_*, *σ_2_* (the standard deviation of the likelihood), **γ**_1_ and **γ**_2_ (perceptual bias). Each parameter can be specified as a free parameter, or can be manually fixed at a value. The GUI’s built-in plotting functions enable users to visualize the fitting results.

The user can choose between two different methods for parameter optimization: a) the ‘Powell’ algorithm from the ‘*minimize*’ function in the *scipy* package (https://scipy.org) or b) the ‘*VBMC*’ method from the *pyvbmc* package (https://acerbilab.github.io/pyvbmc/), which is an approximate Bayesian inference method designed to fit computational models with a limited budget of potentially noisy likelihood evaluations, useful for computationally expensive models or quick inference and model evaluation (Acerbi, 2018; 2020).

#### Model Simulation

The simulation function allows exploring the effect of different parameter values on the behavioral outcome (i.e., response distributions). This will help users gain an intuitive understanding of BCI. It also allows investigation of the behavior of BCI under different parameter or stimulus regimes, which can be useful for qualitative comparison of empirical data with the model. In cases of missing or impoverished data sets that may not allow for reliable data fitting, model simulation can be used to qualitatively compare patterns of empirical data with simulated data. The BCI Toolbox provides five parameters for use:

*p_common_*,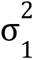 (controlling the variance of modality 1 likelihood), 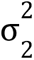 (controlling the variance of modality 2 likelihood) 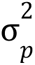 (controlling the variance of prior for perceptual variable of interest) and *μ_P_* (the mean of the prior for perceptual variable of interest). Upon setting these parameters and choosing the value of stimuli (e.g., location of each stimulus in a localization task) to observe, the toolbox generates the simulated data and visualizes it. Researchers can examine subsequent data and compare the perceptual responses under the three decision-making strategies (*model averaging* vs. *model selection* vs. *probability matching*, Figure. 1C). The model simulation module contains one-dimensional continuous data simulation, two-dimensional continuous data simulation and one-dimensional discrete data simulation.

#### Parameter recovery

The reliability of the BCI model was assessed through parameter recovery tests. To evaluate the performance of our model, we first simulate synthetic data with known ground truth parameters. We generated a set of 5 random parameters, where *p_common_*∼ *U*(0, 1), **σ***_1_*∼ *U*(0. 1, 3); **σ***_2_*∼ *U*(0. 1, 3); **σ***_P_*∼ *U*(0. 1, 3); *μ_P_*∼*U*(0, 3. 5). Subsequently, we generated synthetic data using these parameters under the discrete data model structure and through *model averaging*.

Next, we used the BCI toolbox data fitting module to fit the model to the data. Using 10,000 as the *number of simulations*, *minus log likelihood* as the fit type, and *model averaging* as a strategy, we fitted the synthetic data once using the Powell algorithm and once using the VBMC method. Figure. 3 shows the results of parameter optimization for each of these two methods.

**Figure 3.**
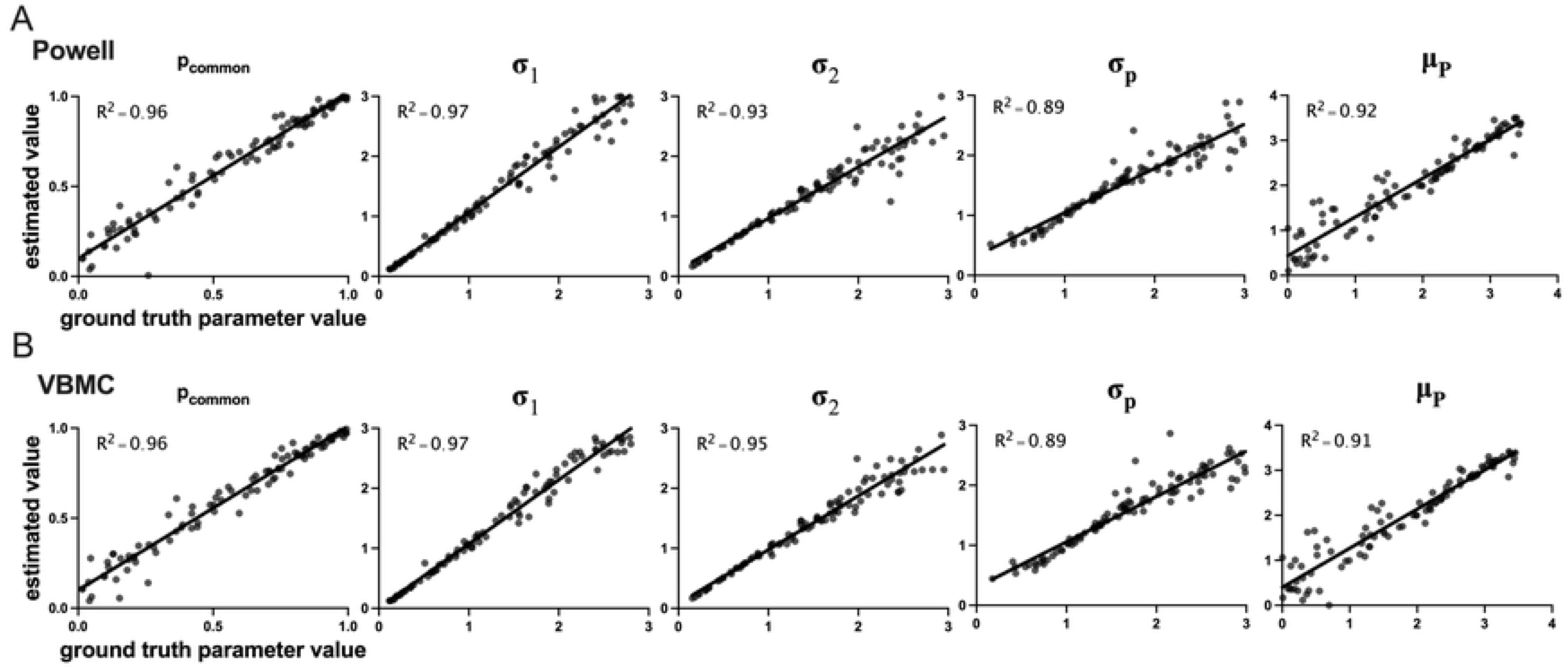
Results of parameter recovery analysis. We generated 100 sets of synthetic data by selecting random values for the 5 model parameters using the discrete 1-dimensional model simulation module of the toolbox. Next, the synthetic data were fitted by the data fitting module of the toolbox. In each panel, the estimated parameter value from data fitting is plotted against the ground-truth value of that parameter. R^2^ indicates the degree of correlation between the estimated and true parameters. In all cases, the model parameters were recovered well. (A) Results from using the Powell algorithm for parameter optimization. (B) Results from using the VBMC method for parameter optimization.

## Results

Here, for the purpose of illustration, an example of the results produced by the toolbox for each of the two main functions–data fitting and model simulation. For the data fitting we used data from Experiment 5 in the study by Odegaard et al. (2017) which is publicly available. In this experiment observers were presented with simple visual and auditory stimuli at one of 5 possible positions along the azimuth, and were asked to report the perceived location of each stimulus in each trial. The responses were provided using a joystick along a continuous horizontal scale on the screen, and therefore were continuous. In that experiment, participants’ spatial localization was tested using a test session as described above. Following the test session, the participants were passively presented with auditory-visual stimuli in an “adaptation” phase. Immediately, after the adaptation phase, the participants were tested again in a spatial localization test identical to the pre-adaptation test. The study reported a statistically-significant increase in *p_common_*after adaptation. We analyzed the behavioral data from the spatial localization tasks using the continuous one-dimensional data fitting module with 5 free parameters (*p_common_*, **σ***_1_*, **σ***_2_*, **σ***_P_* and *μ_P_*), the Powell parameter optimization method, and *model averaging*. Figure 4a shows the results of data fitting for the pre-adaptation test. The toolbox results replicated the finding of Odegaard et al. 2017 study, including the increase in *p_common_*after adaptation (Supplementary File).

**Figure 4.**
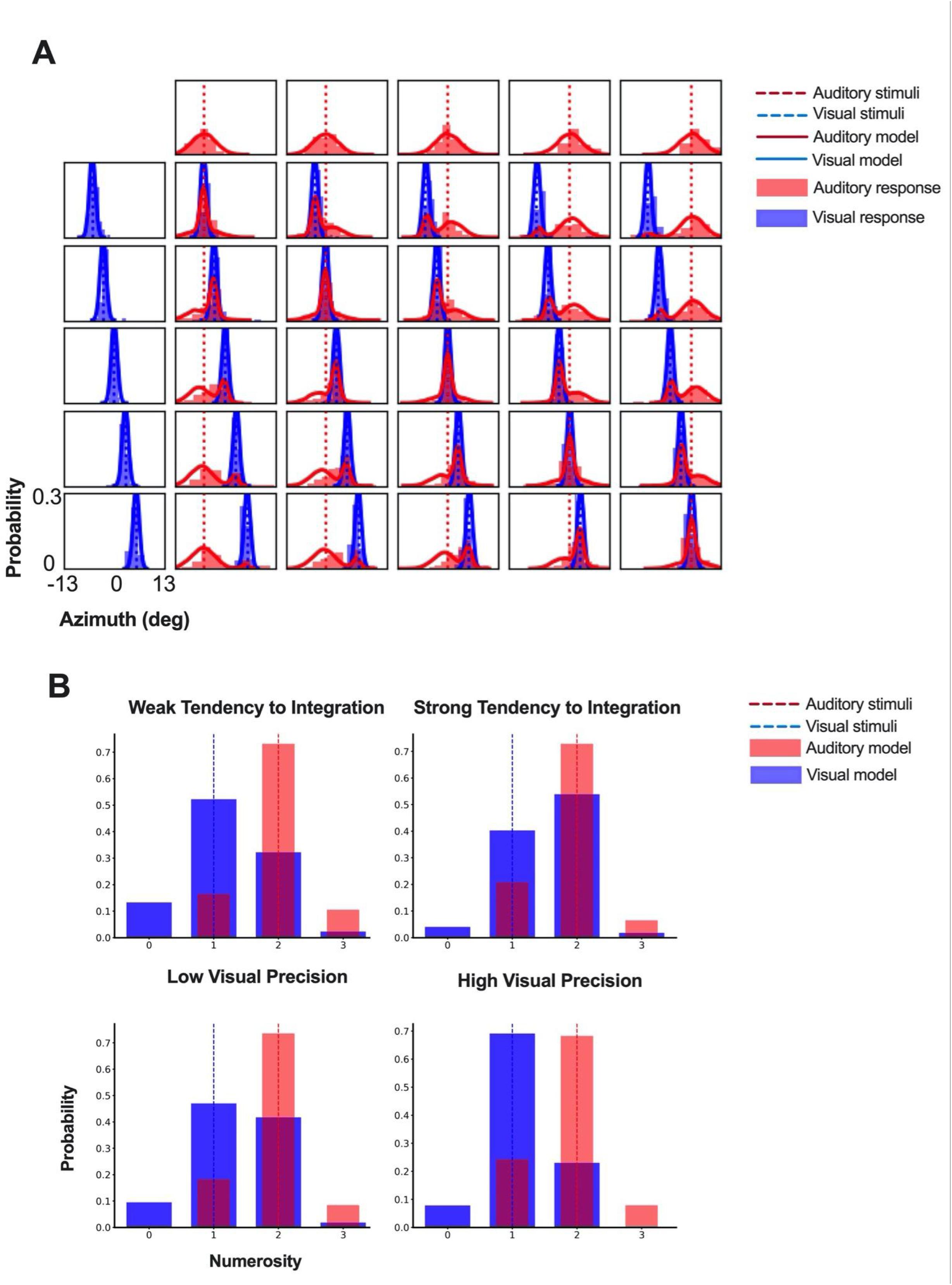
Examples of BCI toolbox outputs. (A) The model fitting results with the continuous data from a spatial localization task. Each plot corresponds to one of the stimulus conditions, with the first row plots representing unisensory auditory conditions (stimulus position varying from left-most to right-most positions along azimuth from left to right), and first column representing unisensory visual conditions, and all other plots corresponding to bisensory conditions. Positions of the auditory and visual stimuli are denoted using broken red and blue vertical lines, respectively. The red and blue histograms represent the auditory and visual response distributions of a specific subject, respectively. The red and blue solid lines represent the model fits produced by the toolbox. (B) The simulation results for one visual stimulus accompanied by two auditory stimuli. We used the fixed parameters (Weak tendency: *p_common_*=0.2; Strong tendency: *p_common_*=0.8; **σ***_1_*=1; **σ***_2_*=0.5; **σ***_P_*=1.5; *μ_P_*=1.5) to simulate how prior integration tendency influences multisensory numerosity perception. We also used the fixed parameters (*p_common_*=0.5; Low visual precision: **σ***_1_*=1; high visual precision: **σ***_1_*=0.5; **σ***_2_*=0.5; **σ***_P_*=1.5; *μ_P_*=1.5) to simulate how unisensory precision influences multisensory numerosity perception.

Figure 4B shows the results of model simulation under 4 different parameter regimes in the one-dimensional discrete module. As an example, we considered the temporal-numerosity task in which the observer task is to report the perceived number of flashes and beeps. The responses in this task are discrete. The number of flashes was set to 1 and the number of beeps was set to 2. In the first row we show the effect of Pcommon on the responses, by keeping all parameters the same but changing the value of Pcommon. The left panel in the top row shows the results for a small Pcommon value (low tendency to integrate), whereas the right panel of the top row shows the results for a large *p_common_*value (high tendency to integrate). Given the higher precision of the auditory modality, the visual perception (perceived number of flashes) is biased by the number of beeps, but the degree of bias is influenced by the value of Pcommon, with a stronger bias (more illusion) observed in the case of higher tendency for integration. The bottom row shows the effect of visual precision (**σ***_V_*) on the responses, by keeping all parameters the same but changing the value of **σ***_V_*. A lower visual precision leads to a stronger bias and a more pronounced illusion in the visual modality. These example simulations illustrate BCI Toolbox’s utility as a tool for gaining an understanding of the model as well as predicting behavioral outcomes in different settings.

## Discussion

We introduced a toolbox for the Bayesian causal inference model that supports the analysis and simulation of behavioral data in a wide range of tasks in multisensory perception and sensorimotor science. BCI toolbox provides modeling tools for a wide range of experimental paradigms and data types and offers a variety of computational and optimization methods under the Bayesian framework. In addition, the BCI toolbox can batch process and visualize the analysis results, promoting understanding and utility of the BCI model.

### Major advantages of BCI Toolbox

One of BCI Toolbox’s primary advantages is its user-friendly GUI, enabling and facilitating the usage of hierarchical Bayesian causal inference models in neuroscience research even by researchers without any computational training. The BCI Toolbox is suitable for new users to learn and utilize the BCI model. The simulation section can be used for pedagogical purposes, as it enables users to intuitively understand the role of various parameters, which can help them further understand the algorithm in the model. Moreover, the simulation functions are useful for qualitative modeling, offering insights into system behaviors beyond mere reliance on quantitative data.

Besides model simulation, the model fitting module is helpful in the quantitative analysis of data, enabling precise parameter estimation and ensuring a more accurate representation of underlying trends behind the behavioral data. The toolbox also offers a model comparison option that enables a thorough comparison of the three decision strategies. Additionally, it supports evaluations based on varying numbers of free parameters.

In addition to the functional advances, we verified the reliability of the BCI model through parameter recovery. The results show that the vast majority of the parameters can be well estimated by the model with an error margin of 5% or less. The current work provides compelling evidence of the scientific validity and reproducibility of the BCI model, offering a reliable data processing option for future cognitive neuroscience research.

### Potential limitations of BCI Toolbox

A notable constraint of the toolbox is the fixed number of variables in each model. Therefore, the users might face challenges in scenarios where flexible configurations or customizations of variables are required, potentially hindering the adaptability of the tool for diverse research applications. Also, although current methods like computing log likelihood or sum of squared errors are used in the BCI Toolbox to measure model-behavioral data discrepancies, improvements are needed. The loss function in the brain is shaped by evolution or experience to minimize specific costs, which varies among individuals and over time (for a review see, Shams & Beierholm, 2022). So we will continue to explore additional possible methods of quantifying the error which may yield better fits. Updates to the BCI Toolbox and its documentation will be provided in due course to reflect these advancements.

In summary, the BCI Toolbox integrates the resources of cognitive neuroscience research that BCI models can interpret in the past decade. It applies the latest algorithms and parameter optimization methods, providing a convenient, reliable, and diverse data processing tool for potential studies. By utilizing top-notch datasets and cutting-edge models, the present work greatly enriches the computing community of cognitive neuroscience. We encourage fellow community members to contribute to its improvement by suggesting improvements, reporting bugs, and offering bug fixes, new ideas, and innovative modifications.

---

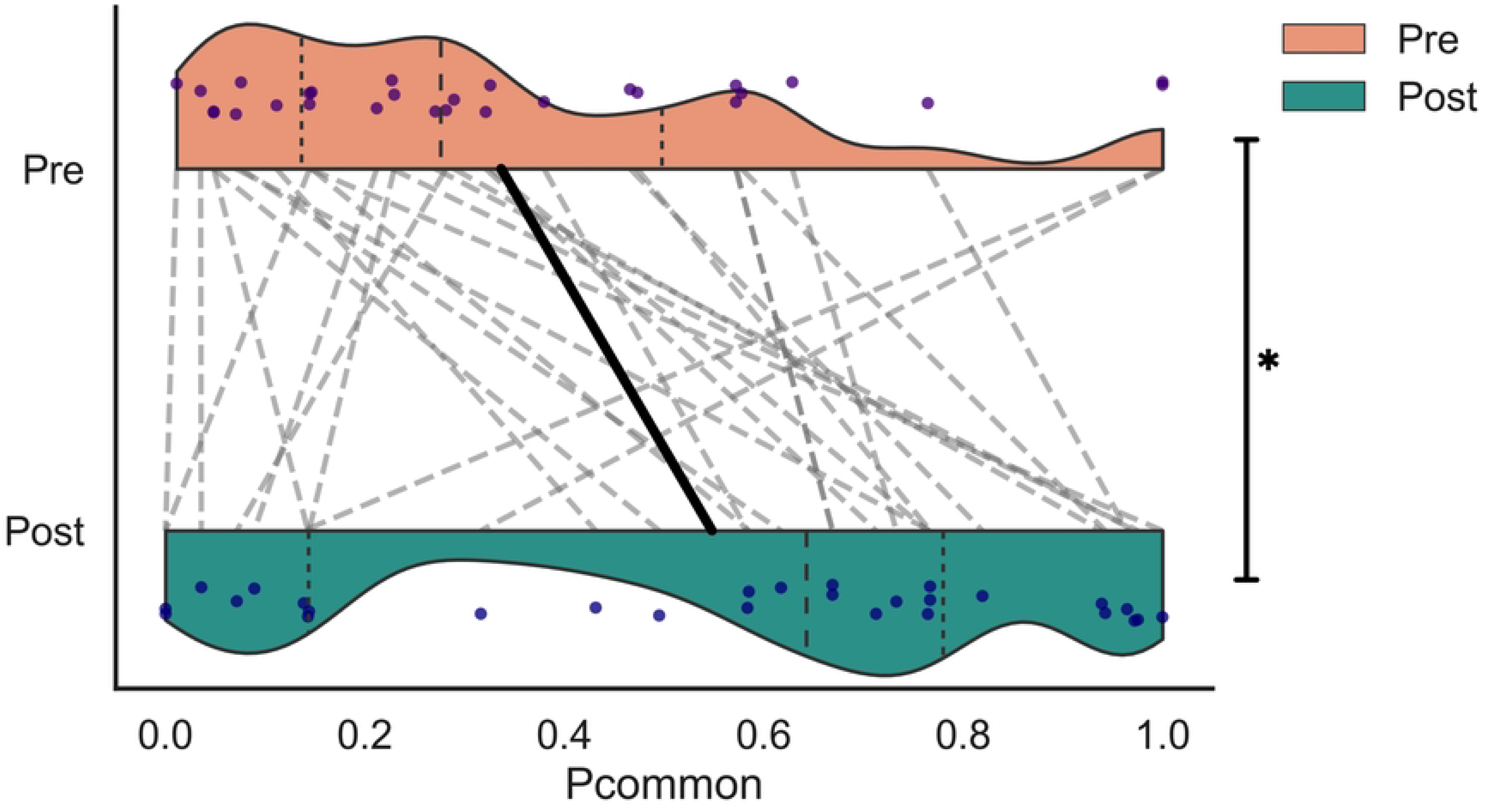

